# Approximate k-nearest neighbors graph for single-cell Hi-C dimensional reduction with MinHash

**DOI:** 10.1101/2020.03.05.978569

**Authors:** Joachim Wolff, Rolf Backofen, Björn Grüning

## Abstract

Single-cell Hi-C interaction matrices are high dimensional and very sparse. To cluster thousands of single-cell Hi-C interaction matrices they are flattened and compiled into one matrix. This matrix can, depending on the resolution, have a few millions or even billions of features and any computation with it is therefore memory demanding. A common approach to reduce the number of features is to compute a nearest neighbors graph. However, the exact euclidean distance computation is in *O*(*n*^2^) and therefore we present an implementation of an approximate nearest neighbors method based on local sensitive hashing running in *O*(*n*). The presented method is able to process a 10kb single-cell Hi-C data set with 2500 cells and needs 53 GB of memory while the exact k-nearest neighbors approach is not computable with 1 TB of memory.

## Introduction

The chromosome conformation capture technique 3C (1) and its successors 4C (2, 3), 5C (4) and Hi-C (5) have given insights over the last years on the spatial proximity of the DNA and its impact on gene regulation. Single-cell Hi-C (6) extends this to individual cell and provides insights into the processes of cell differentiation and division with respect to the dynamics of chromosome confirmation. While Hi-C data analysis demands high computational resources, singlecell Hi-C increases this demand due to the need to not only process one interaction matrix but potentially several thousand. Cell clustering is one of the most important parts of single-cell Hi-C data analysis to gain information about the similarity and therefore the linkage between different cells. To be able to use classical clustering algorithms, scHiCEx-plorer flattens the interaction matrices of a cell to one dimension and creates a new matrix where each row represents one cell. The downside of this approach is a high feature number; for example with 1 megabase (Mb) resolution matrices, and the mice mm9 reference genome, 7.6 million features are present, while using 10 kilobase (kb) matrices the matrix has 76 billion features. The large number of features makes processing single-cell Hi-C data very memory demanding. Moreover, the high memory usage with non-approximate methods makes the clustering inaccessible to a majority of researchers due to the lack of computers with > 1 TB of memory.

## Methods

Computing a k-nearest neighbors graph to reduce the number of features is a well-known approach in single-cell data analysis. With a k-nearest neighbors graph, the number of features is reduced to the number of cells. The exact k-nearest neighbors graph algorithm has a run time of *O*((*d*\**n*)^2^), with *d* the number of dimensions or features and *n* the number of cells. As long as *d* is reasonably small, the computation time will mainly depend on the number of cells *n*, but as the number of features rises to the millions, the compute time becomes more dependent on the features and not on the number of cells. This phenomenon is known as the curse of dimensionality. In single-cell Hi-C the number of features can be in the billions while the matrices are at the same time very sparse. The exact computation of the k-nearest neighbors graph is not only time consuming, but, even more importantly, memory intensive. As shown in Table 1 the memory footprint is a manageable 6.7 GB for a 1 Mb resolution matrix; however, on the 10 kb matrix the memory usage is more than 1 TB. These high requirements are problematic for researchers; access to a computer with a terabyte or more of memory requires either a long waiting time on a shared cluster or is simply not possible. To overcome these limitations, scHiCExplorer provides an approximate k-nearest neighbors graph computation based on MinHash (7), a local sensitive hash function technique.

**Table 1.** Data from (12) Diploid cells, with 7 clusters. For clustering K-means and spectral clustering are used, MinHash with 800 hash functions. All results computed on 2x XEON E5-2630 v4 @ 2.20GHz 2x 10 cores / 2x 20 threads, 1 TB memory.

| Method | Runtime | Memory |
| --- | --- | --- |
| Raw and K-Means | 3:21 h | 33 GB |
| Raw and Spectral | 3:02 min | 6.7 GB |
| PCA and K-Means | 7:27 min | 220 GB |
| PCA and Spectral | 7:39 min | 220 GB |
| sklearn k-nn k = 100 and K-means | 2:11 min | 6.7 GB |
| sklearn k-nn k = 100 and Spectral | 3:40 min | 6.7 GB |
| sklearn k-nn k = 2460 and K-means | 18:41 min | 6.7 GB |
| sklearn k-nn k = 2460 and Spectral | 3:44 min | 6.7 GB |
| MinHash k = 100 and K-means | 2:18 min | 6.7 GB |
| MinHash k = 100 and Spectral | 3:27 min | 6.7 GB |
| MinHash k = 2460 and K-means | 6:59 min | 6.7 GB |
| MinHash k = 2460 and Spectral | 3:20 min | 6.7 GB |
| MinHash exact mode k = 100 and K-means | 4:25 min | 6.7 GB |
| MinHash exact mode k = 100 and Spectral | 2:56 min | 6.7 GB |
| MinHash exact mode k = 2460 and K-means | 1:12 h | 6.7 GB |
| MinHash exact mode k = 2460 and Spectral | 1:03 h | 6.7 GB |

| Method | Runtime | Memory |
| --- | --- | --- |
| Raw and K-Means | - | > 1 TB |
| Raw and Spectral | - | > 1 TB |
| PCA and K-Means | - | > 1 TB |
| PCA and Spectral | - | > 1 TB |
| sklearn k-NN and K-Means | - | > 1 TB |
| sklearn k-NN and Spectral | - | > 1 TB |
| MinHash k = 2508 and K-Means | 1:13 h | 53 GB |
| MinHash k = 2508 and Spectral | 1:04 h | 53 GB |
| MinHash exact mode k = 2508 and K-Means | 3:21 h | 53 GB |
| MinHash exact mode k = 2508 and Spectral | 3:09 h | 53 GB |
| MinHash exact mode k = 50 and K-Mans | 1:10 h | 53 GB |
| MinHash exact mode k = 50 and Spectral | 1:01 h | 53 GB |
| MinHash exact mode k = 200 and K-Mans | 1:19 h | 53 GB |
| MinHash exact mode k = 200 and Spectral | 1:11 h | 53 GB |

(b) Data: 10 kb resolution, with 2460 cells. The raw matrix, PCA and sklearn k-nn methods requested more than the available 1 TB of memory and could not be computed.

### A. MinHash

Local sensitive hash functions are able to hash similar values to an equal data range while not losing the neighborhood relation. This means a local sensitive hashing can be used to compute the similarity between two cells in a very sparse data set. Cells which share features are more likely to be similar to each other compared to cells with less common features. This fact is used by MinhHash (7); for each cell *x*, only the bins ids *x_i_* of non-zero interactions are considered (similar to (8)) and the hash value per hash function is given as:

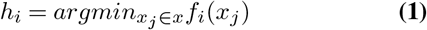

This requires a family of hash functions with the following conditions:

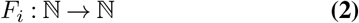

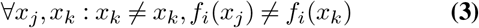

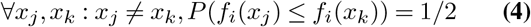

The number of features two cells *x* and *y* share is given by the Jaccard similarity:

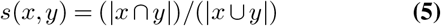

Broder (7) shows that MinHash is a unbiased estimator of the Jaccard similarity:

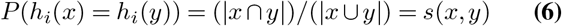

### B. Inverse index

A fast computation of a k-nearest neighbors graph requires a linear run time and a significant reduction of the number of features to overcome the curse of dimensionality. To achieve this, the MinHash algorithm is used to create an inverse index; this inverse index guarantees a reduction of query dimensions to the number of used hash functions of MinHash. However, an additional linear fitting in contrast to method is required.

#### B.1. Fitting

To reduce the number of dimensions in linear runtime, MinHash considers for one hash function only the non-zero ids of a cell, as seen in Equation 1. All MinHash values of all hash functions together are called the *signature* of cell. These signatures are inserted to an inverse index to achieve a fast query time. The inverse index stores for each hash function the hash values and in which cell they were occurring. With this approach it is possible to reduce the number of features to the number of hash functions per cell id. The runtime of the fitting depends on the number of cells *n*, the number of hash functions *h* and the number of non-zero features per cell *f* and is given as *O*(*n*\**h*\**f*). For an example of the fitting and the inverse index structure, please consider Figure 1.

**Fig. 1.**
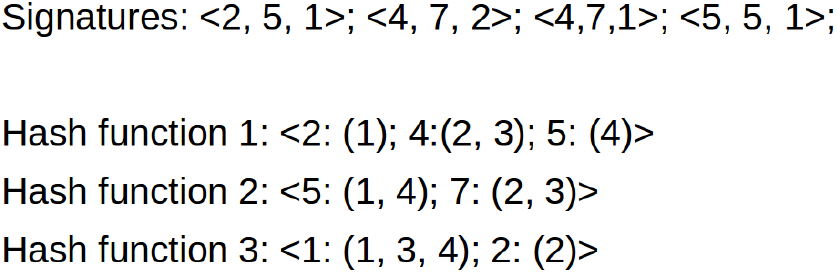
An exemplary signature and inverse index: The signature is created for four cells and three hash functions. The inverse index stores for each hash function the computed hash value and the id of the cell. For example, for the second cell < 4, 7, 2 > the first hash function *Hash function* 1 stores the computed hash value *4* and associates the id of the cell: *Hash function* 1 :< 2 : (1); **4** : (**2, 3**); 5 : (4) >. The same hash function and hash value occurs for cell number three again, this is a collision of hash function 1 for cell 2 and cell 3.

#### B.2. Collision based approximate nearest neighbors graph

The number of hash collisions between two instances gives an estimate of their similarity, as seen in Equation 6. To achieve this, the signature of a cell is used to search for *hash collisions* in the inverse index. A *hash collision* between two cells is defined as the same hash value for the same hash function. The more collisions two cells have, the closer they are. The query time of this approach is depending only on the number of used hash functions and, if not stored in memory from the fitting phase, the computation of signatures. Additional a negligible factor caused by the sorting of all occurrences of collisions and the query time of the used data structures of the inverse index, must be considered for the run time.

#### B.3. Euclidean distance

MinHash is an estimator for the Jaccard similarity but not for the euclidean distance which is an often used measurement for similarity of data and therefore for a k-nearest neighbors graph. To increase the accuracy of the computed k-nearest neighbors graph with respect to the euclidean distance, the option to use an exact mode is given. First, the nearest neighbors based on MinHash to reduce the search space is computed. To achieve a higher accuracy, the nearest neighbors of a cell and also the neighbors of the neighbors are computed. This strategy is a result of the observation that MinHash misses sometimes a true neighbor, but a neighbor of a neighbor is detect. This subset of cells is used as an input to compute the exact nearest neighbors using the euclidean distance. See Figure 2.

**Fig. 2.**
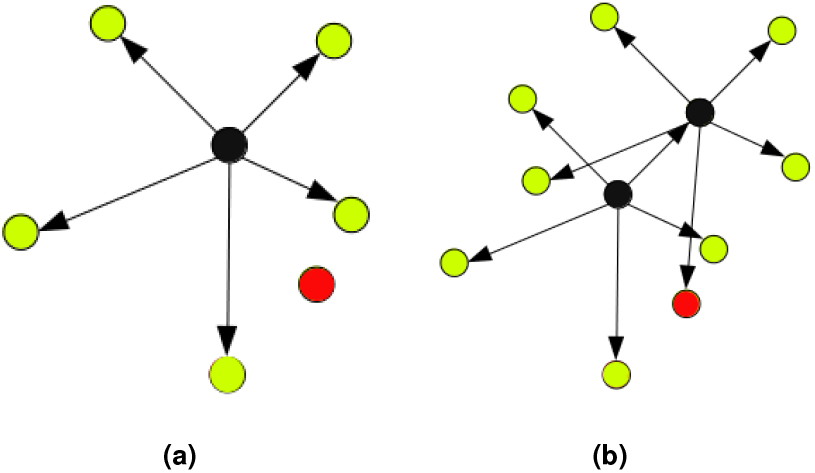
Detected neighbors with MinHash. **(a)**: The red node is close to the black node, but MinHash can fail to detect it due to its approximate nature. However, **(b)** shows it is likely that a neighbor of the node has the red node in its neighborhood. For this reason, the neighbors of the neighbors are considered in the slower exact mode mode too.

### C. Implementation

To compute the approximate nearest neighbors with MinHash, a highly optimized library ‘sparse-neighbors-search’ was implemented in C++ with SSE and OpenMP support. Hash values may optionally be computed with general purpose GPU (GPGPU) processing with Nvidias CUDA. To ensure user accessibility, the C++ library is embedded to a Python 3.6 interface. The MinHash approximation of a k-nearest neighbors graph is part into the scHiC-Explorer, which can handle all pre-processing steps. These include the demultiplexing of raw FASTQ data, quality control, the merging of the interaction matrix per cell to one scool matrix (9), the read coverage normalization, correction with the KR algorithm (10) and methods for scool matrix manipulations. Subsequent to the dimension reduction, the clustering based on the approximate k-nearest neighbors graph can be applied with k-means or spectral clustering. An option to visualize the clustering result as a consensus matrix or matrix profiles is given. Besides its embedding in the scHiC-Explorer, the approximate nearest neighbors implementation can be used by any Python 3.6 software, as its API is similar to the nearest neighbors API of sklearn (11).

The benefits of the MinHash technique are present at many levels. The approximate Jaccard similarity computation achieves faster or similar run times in comparison to the sklearn implementation of a k-nearest neighbors search, which is based on ball-trees; under consideration of the one megabase resolution single-cell Hi-C dataset as shown in Table 1a. The runtimes are heavily influenced by the used clustering algorithm, K-means runs usually slower. This is especially the case for the clustering on raw data, K-means runs for more than three hours, for all others a difference is present, but acceptable because it is in the range of minutes. The classical and naive way to reduce dimensions is a principal component analysis (PCA), but the method uses an outstanding amount of memory (220 GB) already on the low resolution matrix. All k-nearest neighbors graph approaches are using the same amount of memory which is caused by the memory consumption at the read-in stage of the individual single-cell Hi-C matrices. The provided exact mode of MinHash has similar run times in comparison to sklearns implementation, however it is outperformed if it has to compute the full nearest neighbors graph (18 min vs 1:12 h). The benefits of MinHash in terms of runtime and memory usage are more significant under the consideration of a high resolution single-cell Hi-C data set. As shown in Table 1b all methods beside MinHash are not able to be computed caused by their high memory usage of more than one terabyte. Only the MinHash approach can compute a result while it is using moderate 50 GB of memory; resources requirements which are available for the most researchers.

The major share of the runtime of the presented clustering approaches is caused by the load process of the individual single-cell Hi-C matrices into the memory. For example, to load the 10 kB Hi-C matrices with 40 threads takes around 58 minutes, while the fitting and approximate k-nearest neighbors graph is computed within a few seconds, see Table 2a and 2b. However, this is different for the full nearest neighbors graph. To compute all euclidean distances of the neighbors and their neighbors as shown in Section B.3 takes time. The important result is that the full nearest neighbor matrix can be computed with a reasonable amount of memory. Considering the time for the read in of the matrices, the time that is required to hash all non-zero ids and to create the inverse index is minor. The hashing time with 800 hash functions was measured with 20 cores / 40 threads and compared to the time a Nvidia Titan T4 needed. The runtime on the GPU was in total one minute less. This result shows the usage of GPGPU can improve the performance a bit, but is negligible.

### D. Accuracy

The chromatin structure of cells of the same cell stage are naturally quite equal. A valid distinction is often not possible and an accuracy measure based on the rank of the detected nearest neighbors is potentially misleading. For this reason, the used accuracy measure is defined as the intersection between the nearest neighbors of a cell as detected by the exact scikit-learn implementation and the results by MinHash respectively the MinHash exact mode implementation.

Let *X* = {*x*_0_,…, *x_k_*} be an approximate k-nearest neighborhood and *Y* = {*y*_0_, …, *y_n_*} an exact k-nearest neighborhood. The accuracy by intersection is defined as:

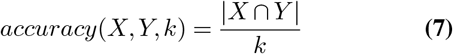

Let *A* = {*a*_0_, …, *a_n_*}, |*A*| = *n* the approximate k-nearest neighbors and *B* = {*b*_0_, …, *b_n_*}, |*B*| = *n* the exact k-nearest neighbors; where *n* is the number of cells. The accuracy for these two k-nearest neighbors graphs is given as:

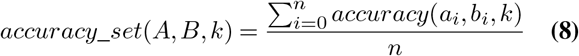

The accuracy of approximate algorithms is crucial to prove that the run time and memory benefits are not caused by larger inaccuracies. The here presented MinHash algorithm is compared to the results of sklearn’s k-nearest neighbor graph which is based on the euclidean distance. It must be noted that MinHash itself is an approximation of the Jaccard similarity and not of the euclidean distance. This becomes obvious considering the results for the MinHash algorithm in Table 2. The exact mode of MinHash computes the euclidean distances on a pre-selection of neighbors and is therefore an approximate algorithm for an euclidean k-nearest neighbors graph. This approach achieves very high accuracies. The 50-nearest neighbors graph achieves an accuracy of 0.97 for the 1 MB resolution matrix and 0.81 for the 10 kb resolution matrix. Considering the 100-nearest neighbors graph, an accuracy of 0.98 respectively 0.92 can be reached; for250-nearest neighbors and more, an accuracy greater 0.99 is given.

**Table 2.**
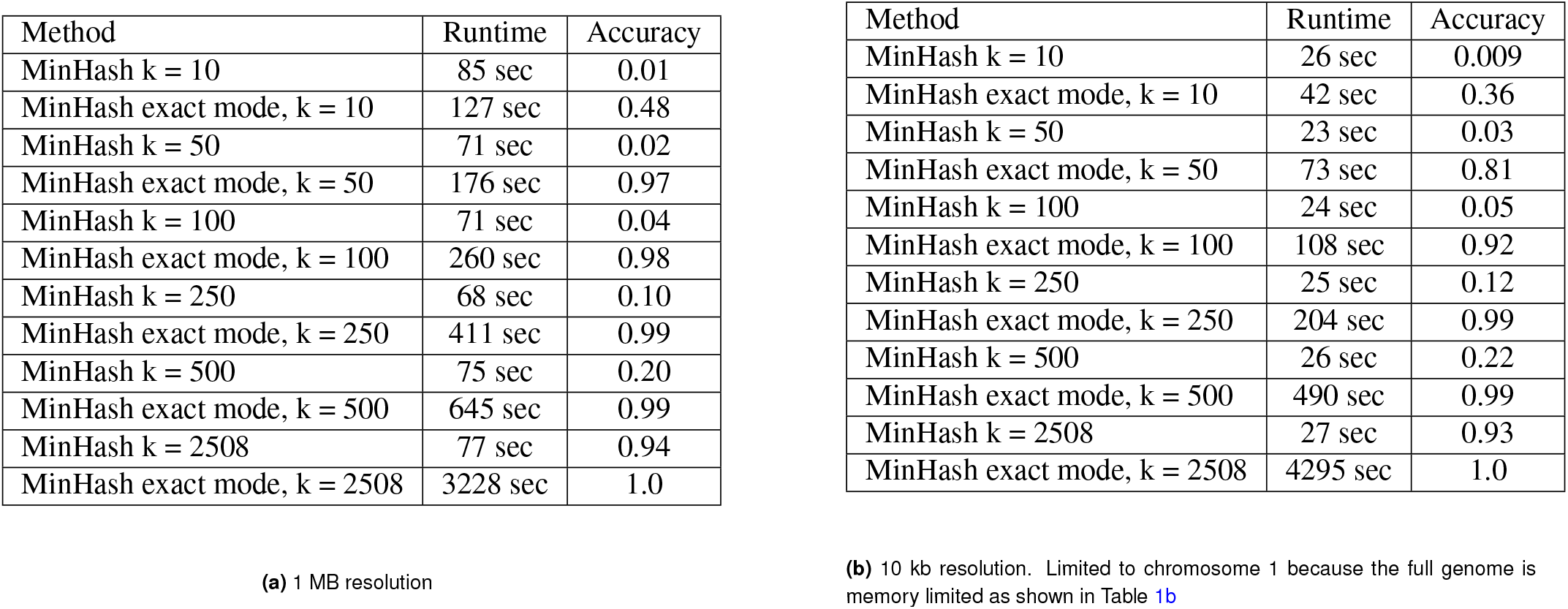
Accuracy measure of k-nearest neighbors. Comparison of sklearn k-nearest neighbors results which are based on euclidean distance with MinHash and MinHash exact mode. It clearly shows that MinHash is an estimator of the Jaccard similarity but not of the euclidean distance. The results can be used to decrease the search space in the exact mode. All data from (12), with 2508 cells. The run times are without the loading of the individual files and directly on the compiled, usually intermediate, raw scHi-C matrix. The used number of hash functions for MinHash is 800. All results computed on XEON E5-2630 v4 @ 2.20GHz 10 cores/ 20 threads, 120 GB memory.

The three methods to reduce the number of dimensions via k-nearest neighbors graph have a different impact on the results of the clustering. Figure 3 shows the results of the clustering applied on either a 100-nearest neighbors graph or a full nearest neighbors graph; the clustering itself is computed by K-means and spectral clustering. The results confirm the accuracy measures, for example the clustering based on the euclidean 100-nearest neighbors and full-nearest neighbors graph is quite similar between sklearn and MinHash in exact mode, see Figure 3a, 3i and Figure 3c, 3k. The output of spectral clustering shows a different case: The difference between the exact euclidean distance and the approximated MinHash is large, see Figure 3b, 3j and Figure 3d, 3l. It must be noted that the results of MinHash are more accurate and similar to K-Means, see Figure 3i, 3j and Figure 3k, 3l. Also the approximate Jaccard similarity as computed by Min-Hash provides good results and shows it is compatible with the k-nearest neighbors provided via the euclidean distance, see Figure 3e, 3f and 3g. Moreover, it adds an additional option for the clustering process and can, as it is the case for spectral clustering, provide a base for a better result.

**Fig. 3.**
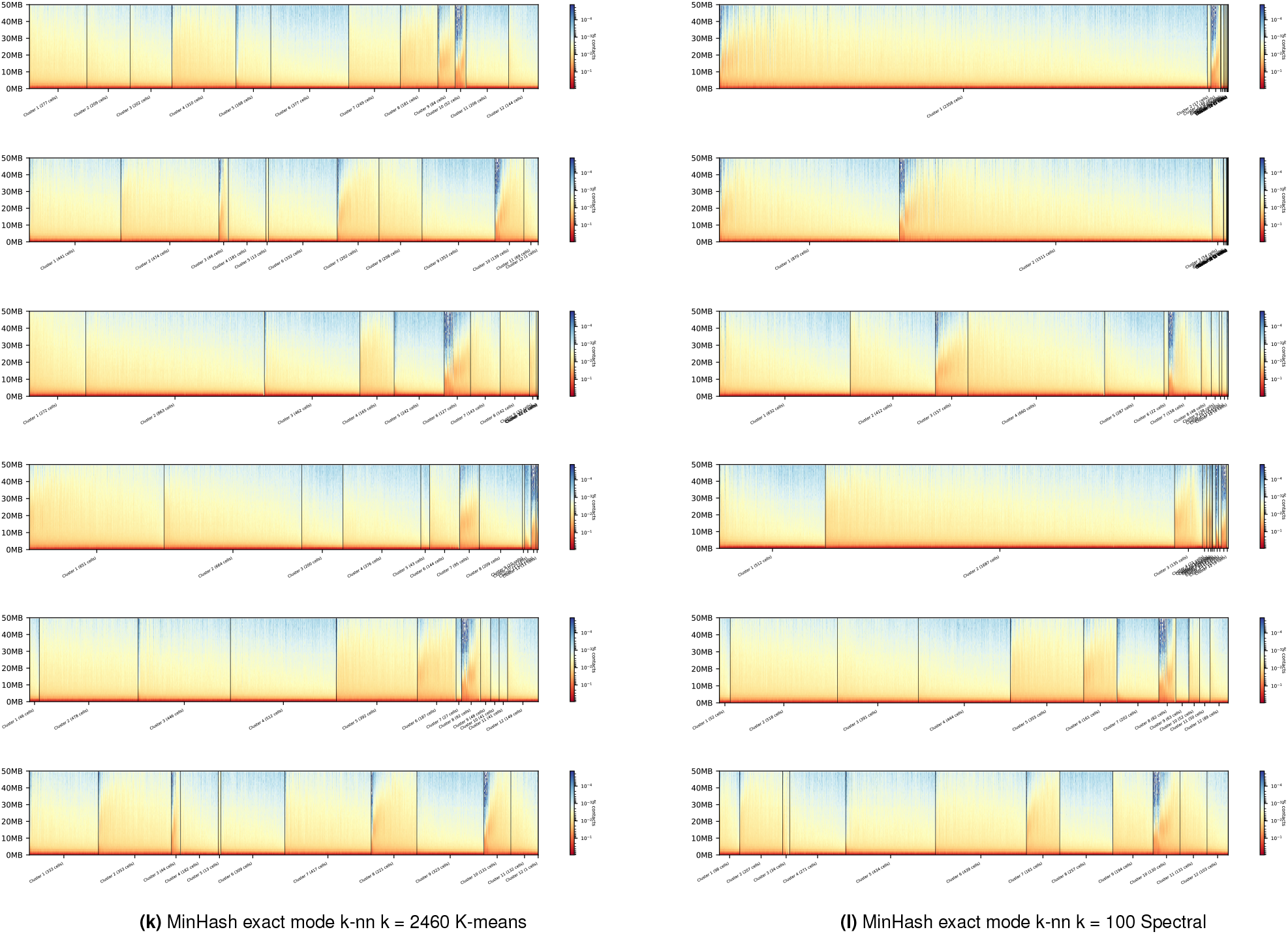
Cluster profiles of K-means and spectral clustering results on 1 MB resolution, with 2460 cells from (12) Diploid cells, with 12 clusters. The cluster profile shows the decreasing contact probabilities of all interactions per cluster member in a genomic distance up to 50 MB. The plots indicate the result depends on the used dimension reduction approach and the clustering algorithm.

## Discussion

The presented approach to reduce the amount of features, especially when dealing with millions to billions of dimensions is an crucial step to achieve acceptable run time and memory usages. More importantly, it is possible to compute a clustering and not have an out of memory exception. Access to computers with more than 1 TB of memory is currently difficult, but an access to computers or cluster nodes with 60 GB of memory is available to the most researchers. The presented approximate nearest neighbor graph enables a broader range of researchers to work with single-cell Hi-C data and adds with the approximate Jaccard similarity another method to create a k-nearest neighbor graph. This method can be useful under the consideration of the problematic nature of different read coverages of the individual single-cell Hi-C matrices. MinHash considers only the presence of non-zero values and is therefore immune to effects based on a different coverage. Moreover, MinHash is embedded into the scHiC-Explorer, a software suite for single-cell Hi-C data analyses. The scHiCExplorer supports the full pipeline of single-cell Hi-C data analysis, starting with demultiplexing, interaction matrix creation, correction, normalization, interaction matrices manipulation, clustering with dimension reduction and visualization. All this is implemented in a highly parallel approach to fully support modern multi-core platforms. This is possible with the use of to the standard cooler file format which enables loading and store multiple matrices in a parallel way. The usage of the multi-cooler format is crucial to store single-cell Hi-C data in a shareable format. With this users can use the raw scool interaction matrix and do not need to deal with the large overhead of data. The two offered modes to compute the approximate nearest neighbors graph shows that the reduction of the search space with Min-Hash and the additional computation of the euclidean distances leads to a k-nearest neighbors graph of high accuracy. The fast mode of MinHash by only using the number of collisions as an approximation of the Jaccard similarity offers an additional k-nearest neighbors graph. This method is, as the accuracies in comparison to the euclidean distance indicate, different, but as it can be seen in Figure 3, can lead to better clustering results. By the availability of MinHash as an independent software package, it can be easily integrated to other research issues dealing with similar matrix properties like it is the case in single-cell RNA-seq.

## Acknowledgements

We thank Simon Bray and Anup Kumar for proof reading the manuscript. We thank Milad Miladi and Fabrizio Costa for the supervision of the implementation of MinHash.

## Bibliography

1 Job Dekker, Karsten Rippe, Martijn Dekker, and Nancy Kleckner. Capturing chromosome conformation. science, 295(5558):1306–1311, 2002. [PubMed:11847345][doi:10.1126/science.1067799].

2. Marieke Simonis, Petra Klous, Erik Splinter, Yuri Moshkin, Rob Willemsen, Elzo De Wit, Bas Van Steensel, and Wouter De Laat. Nuclear organization of active and inactive chromatin domains uncovered by chromosome conformation capture–on-chip (4c). Nature genetics, 38(11):1348, 2006. [PubMed:17033623] [doi:10.1038/ng1896].

3. Zhihu Zhao, Gholamreza Tavoosidana, Mikael Sjölinder, Anita Göndör, Piero Mariano, Sha Wang, Chandrasekhar Kanduri, Magda Lezcano, Kuljeet Singh Sandhu, Umashankar Singh, et al. Circular chromosome conformation capture (4c) uncovers extensive networks of epigenetically regulated intra-and interchromosomal interactions. Nature genetics, 38 (11):1341, 2006. [PubMed:17033624] [doi:10.1038/ng1891].

4. Josée Dostie, Todd A Richmond, Ramy A Arnaout, Rebecca R Selzer, William L Lee, Tracey A Honan, Eric D Rubio, Anton Krumm, Justin Lamb, Chad Nusbaum, et al. Chromosome conformation capture carbon copy (5c): a massively parallel solution for mapping interactions between genomic elements. Genome research, 16(10):1299–1309, 2006. [PubMed:16954542] [PubMed Central:PMC1581439] [doi:10.1101/gr.5571506].

5. Erez Lieberman-Aiden, Nynke L. Van Berkum, Louise Williams, Maxim Imakaev, Tobias Ragoczy, Agnes Telling, Ido Amit, Bryan R. Lajoie, Peter J. Sabo, Michael O. Dorschner, Richard Sandstrom, Bradley Bernstein, M. A. Bender, Mark Groudine, Andreas Gnirke, John Stamatoyannopoulos, Leonid A. Mirny, Eric S. Lander, and Job Dekker. Comprehensive mapping of long-range interactions reveals folding principles of the human genome. Science, 326(5950):289–293, oct 2009. ISSN 00368075. doi: 10.1126/science.1181369. [PubMed:19815776] [PubMed Central:PMC2858594] [doi:10.1126/science.1181369].

6. Takashi Nagano, Yaniv Lubling, Tim J Stevens, Stefan Schoenfelder, Eitan Yaffe, Wendy Dean, Ernest D Laue, Amos Tanay, and Peter Fraser. Single-cell hi-c reveals cell-to-cell variability in chromosome structure. Nature, 502(7469):59, 2013. [PubMed:24067610] [PubMed Central:PMC3869051] [doi:10.1038/nature12593].

7. Andrei Z Broder. On the resemblance and containment of documents. In Proceedings. Compression and Complexity of SEQUENCES 1997 (Cat. No. 97TB100171), pages 21–29. IEEE, 1997. [doi:10.1109/SEQUEN.1997.666900].

8. Steffen Heyne, Fabrizio Costa, Dominic Rose, and Rolf Backofen. Graphclust: alignment-free structural clustering of local rna secondary structures. Bioinformatics, 28(12):i224–i232, 2012.

9. Nezar Abdennur and Leonid A Mirny. Cooler: scalable storage for Hi-C data and other genomically labeled arrays. Bioinformatics, 36(1):311–316, 07 2019. ISSN 1367-4803. doi: 10.1093/bioinformatics/btz540. [PubMed:31290943] [doi:10.1093/bioinformatics/btz540].

10. Philip A Knight and Daniel Ruiz. A fast algorithm for matrix balancing. IMA Journal of Numerical Analysis, 33(3):1029–1047, 2013. [doi:10.1093/imanum/drs019].

11. Lars Buitinck, Gilles Louppe, Mathieu Blondel, Fabian Pedregosa, Andreas Mueller, Olivier Grisel, Vlad Niculae, Peter Prettenhofer, Alexandre Gramfort, Jaques Grobler, Robert Layton, Jake VanderPlas, Arnaud Joly, Brian Holt, and Gaël Varoquaux. API design for machine learning software: experiences from the scikit-learn project. In ECML PKDD Workshop: Languages for Data Mining and Machine Learning, pages 108–122, 2013.

12. Takashi Nagano, Yaniv Lubling, Csilla Várnai, Carmel Dudley, Wing Leung, Yael Baran, Netta Mendelson Cohen, Steven Wingett, Peter Fraser, and Amos Tanay. Cell-cycle dynamics of chromosomal organization at single-cell resolution. Nature, 547(7661):61, 2017. [PubMed:28682332] [PubMed Central:PMC5567812] [doi:10.1038/nature23001].

